# Decentralized multi-site VBM analysis during adolescence shows structural changes linked to age, body mass index, and smoking: A COINSTAC analysis

**DOI:** 10.1101/846386

**Authors:** Harshvardhan Gazula, Bharath Holla, Zuo Zhang, Jiayuan Xu, Eric Verner, Ross Kelly, Gunter Schumann, Vince D. Calhoun

## Abstract

In the recent past, there has been an upward trend in developing frameworks that enable neuroimaging researchers to address challenging questions by leveraging data across multiple sites all over the world. One such framework, Collaborative Informatics and Neuroimaging Suite Toolkit for Anonymous Computation (COINSTAC), provides a platform to analyze neuroimaging data stored locally across multiple organizations without the need for pooling the data at any point during the analysis. In this paper, we perform a decentralized voxel-based morphometry analysis of structural magnetic resonance imaging data across two different sites to understand the structural changes in the brain as linked to age, body mass index and smoking. Results produced by the decentralized analysis are contrasted with similar findings in literature. This work showcases the potential benefits of performing multi-voxel and multivariate analyses of large-scale neuroimaging data located at multiple sites.

## 1 Introduction

Collecting neuroimaging data is expensive as well as time consuming [Landis et al., 2016]. Other significant challenges include the storage and computational resources needed which could prove costly as the volume of the data collected goes up. On the contrary, aggregating information from data across various sites not only makes the predictions more certain by increasing the sample size [Button et al., 2013], but also ensures reliability and validity of the results, and safeguards against data fabrication and falsification [Tenopir et al., 2011, Ming et al., 2017]. In the past few years, there has been a proliferation of efforts [Poldrack et al., 2013] towards enabling researchers to leverage data across multiple sites.

Plis et al. [Plis et al., 2016], proposed a web-based framework titled Collaborative Informatics and Neuroimaging Suite Toolkit for Anonymous Computation (COINSTAC) to such collaborative analysis of data from different sites. COINSTAC implements a privacy-preserving message passing infrastructure that allows large scale analysis of decentralized data. The results thus obtained are on par with those that would have been obtained if the data were in one place. Since, there is no pooling of data it also preserves the privacy of individual datasets.

One such decentralized analysis method available in the COINSTAC framework is voxel-based morphometry [Ashburner and Friston, 2000] which was already discussed in [Gazula et al., 2018] by conceptualizing some variants of the decentralized regression and validating on some publicly available dataset. In this paper, we showcase the power of COINSTAC framework by conducting a real-world decentralized VBM analysis of MRI data at two different sites to study structural changes in adolescent brain as linked to age, body mass index (BMI), and smoking.

Smoking is a major public health concern and an economic burden. It is well known that most adult smokers take up smoking during their adolescent years [Lydon et al., 2014]. However, very little is understood about the brain mechanisms that influence smoking behavior. It is important to understand the effects of smoking on cortical thickness or volume. Understanding the complex neural processes underlying smoking and smoking-induced neural change could be critical to designing interventions to discourage such smoking behavior in adolescents [Ewing et al., 2016].

On the other hand, neuroimaging is becoming increasingly common in obesity research as investigators try to understand the neurological underpinnings of appetite and body weight in humans [Carnell et al., 2012]. This because a higher body mass index (BMI) is associated with structural brain changes, cognitive decline, and an increased risk of Alzheimer disease (AD) in late life [Cronk et al., 2010]. However, there is reason to suspect that the relationship between excess weight and structural brain differences is not limited to older adults [Gunstad et al., 2008].

Our contributions in this paper can thus be summarized as follows.

1. Showcasing decentralized voxel based morphometry on large datasets across multiple sites, IMAGEN from UK and cVEDA from India (more on this later), in the COINSTAC framework and some observations.
2. An understanding of the effects of smoking and body mass index on the structural changes in the brain with age via decentralized voxel-based morphometry.

The outline of the current paper is as follows: In section 2, we discuss the decentralized regression algorithm which forms the bedrock for the decentralized voxel based morphometry analysis followed by an overview of the COINSTAC framework. In section 3, we discuss the IMAGEN and cVEDA data used in this study. In sections 4 and 5, we present the results and discuss our whole experience with using the COINSTAC framework as well as what we found analyzing the results. We conclude with some final comments in section 6.

## 2 Methods

### 2.1 Decentralized VBM (i.e. voxelwise decentralized regression)

Voxel-based morphometry (VBM) [Ashburner and Friston, 2000] is a statistical method that facilitates a comprehensive comparison, via generalized linear modeling, of voxel-wise gray matter concentration between different groups, for example. To enable such statistical assessment on data present at various sites, it is important to develop decentralized tools.

The goal of decentralized regression (the building block of decentralized VBM) is to fit a linear equation (given by equation 1) relating the covariates at *S* different sites to the corresponding responses. Assume each site *j* has data set 𝒟_*j*_ = {(**x**_*i*_, *y*_*i*_): *i* ∈ {1, 2, …, *s*_*j*_}} where **x**_*i,j*_ ∈ *R*^*d*^ is a *d*-dimensional vector of real-values features, and *y*_*j*_ ∈ is a response. We consider fitting the model in equation 2 where **w** is given as [**w**; *b*] and **x** as [**x**; 1]

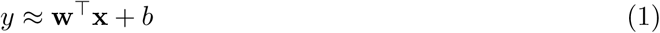

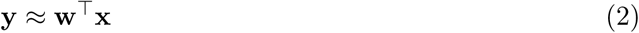

The vector of regression parameters/weights **w** is found by minimizing the sum of the squared error given in equation 3

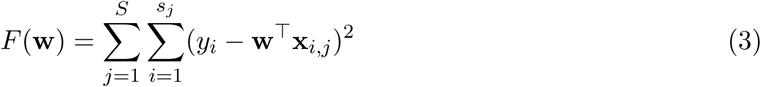

The regression objective function is a linearly separable function, that can be written as sum of a local objective function calculated at each local site as follows:

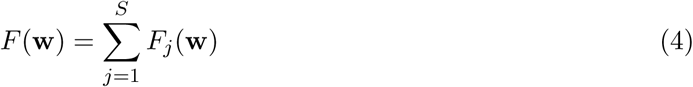

where

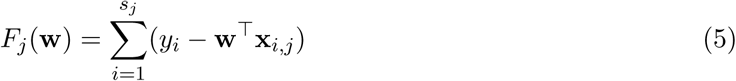

A central aggregator (*AGG*) computes the global minimizer **ŵ** of *F* (**w**).

#### 2.1.1 Decentralized Regression with Normal Equation

For a standard regression problem of the form given in equation 2, the analytical solution is given as

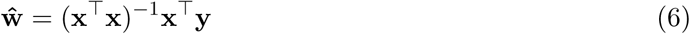

Assuming that the augmented data matrix **x** is made up of data from different local sites, i.e.

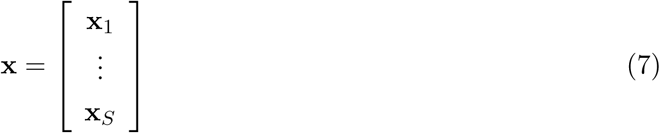

it’s easy to see that **ŵ** can be written as

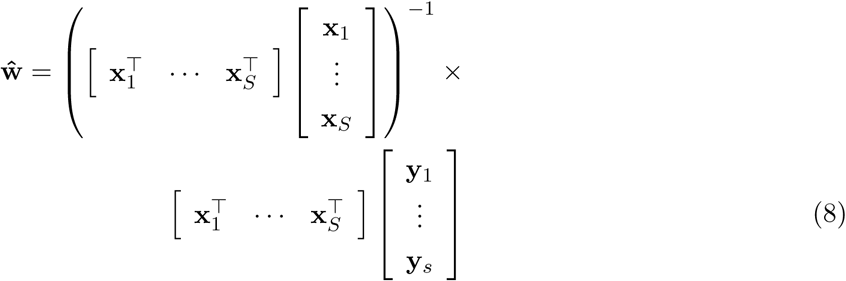

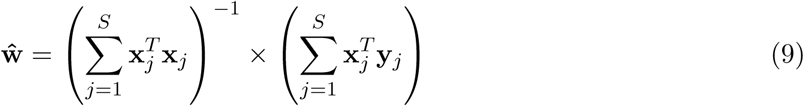

The above variant of the analytical solution to a regression model shows that even if the data resides in different locations, fitting a global model in the presence of site covariates delivers results that are exactly similar to the pooled case.

##### Algorithm 1 Decentralized Regression with Normal Equation

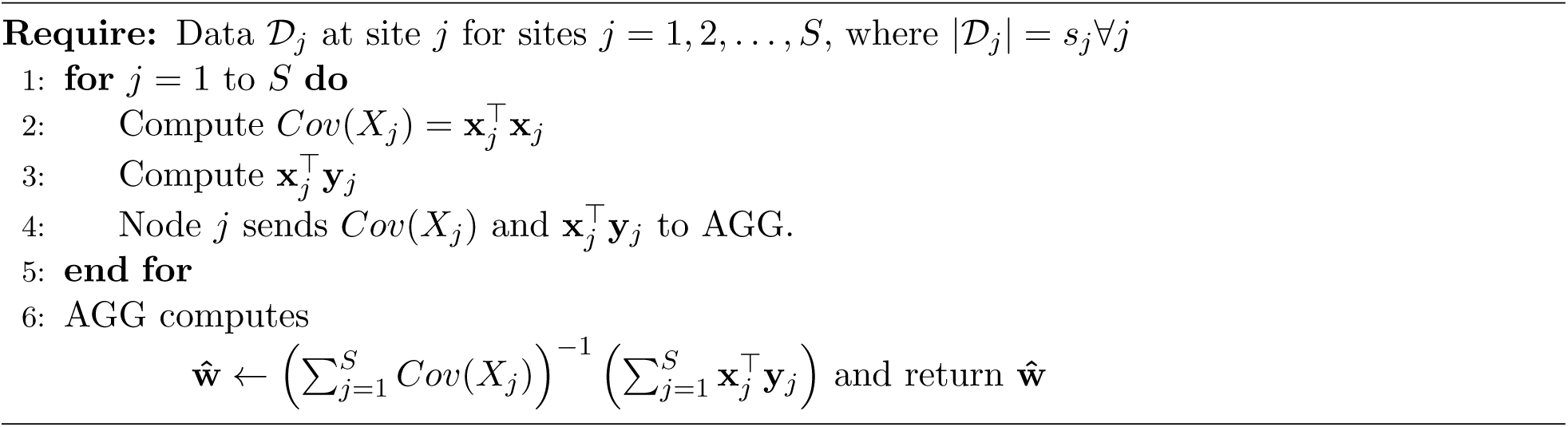

#### 2.1.2 Other Statistics

In addition to generating the weights of the covariates (regression parameters), one would also be interested in determining the overall model performance given by goodness-of-fit or the coefficient of determination (*R*^2^) as well as the statistical significance of each weight parameter (*t* -value or *p*-value).

Determining *R*^2^ entails calculating the sum-square-of-errors (*SSE*) as well as total sum of squares (*SST*) which are evaluated at each local site and then aggregated at the global site to evaluate *R*^2^ given by 1 − *SSE/SST*. An intermediary step before the calculation of *SST* is the calculation of the global 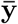 which is determined by taking a weighted average of the local 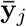 weighted on the size of data at each local site.

The steps involved in calculating the *t* -values (and therefore *p*-values) of each regression parameter are explained in [Gazula et al., 2018, Ming et al., 2017]. The global weight vector (**ŵ**) is sent to each of the local sites where the local covariance matrix as well as the sum-square-of-errors is calculated and sent back along with the data size to the aggregator (*AGG*) which then utilizes that information to calculate the *t* -values for each parameter (or coefficient). Once, the *t* -values have been calculated, the corresponding two-tailed *p*-values can be deduced using any publicly available *distributions* library.

### 2.2 COINSTAC

#### 2.2.1 Description

COINSTAC is an easy-to-install, standalone application with a user-friendly, simple, and intuitive interface. It is open source and freely available for download from https://github.com/ trendscenter/coinstac. It is compatible with Windows, macOS, and Linux operating systems. The software utilizes docker containers (https://www.docker.com/) to run computations. Example of some computations currently available in COINSTAC are independent vector analysis (IVA) [Wojtalewicz et al., 2017], neural networks [Lewis et al., 2020, Lewis et al., 2017], decentralized stochastic neighbor embedding (dSNE) [Saha et al., 2017], joint independent component analysis (ICA) [Baker et al., 2015], and two-level differentially private support vector machine (SVM) classification [Sarwate et al., 2014].

#### 2.2.2 Implementation

The first step to collaborating a decentralized analysis on the COINSTAC platform is creating an account (Figure 1) followed by creating a consortium. A consortium is a group of users who run a decentralized analysis together. Each member/site who’s contributing data for a study create a login and all such members constitute a consortium. There’s one consortium owner who creates the computation. Figure 2 shows an example of the COINSTAC consortium page where there is a card for each consortium containing its name, description, active pipeline, list of owner(s) and members, a button to view details about the consortium, and a button to join the consortium.

**Figure 1:**
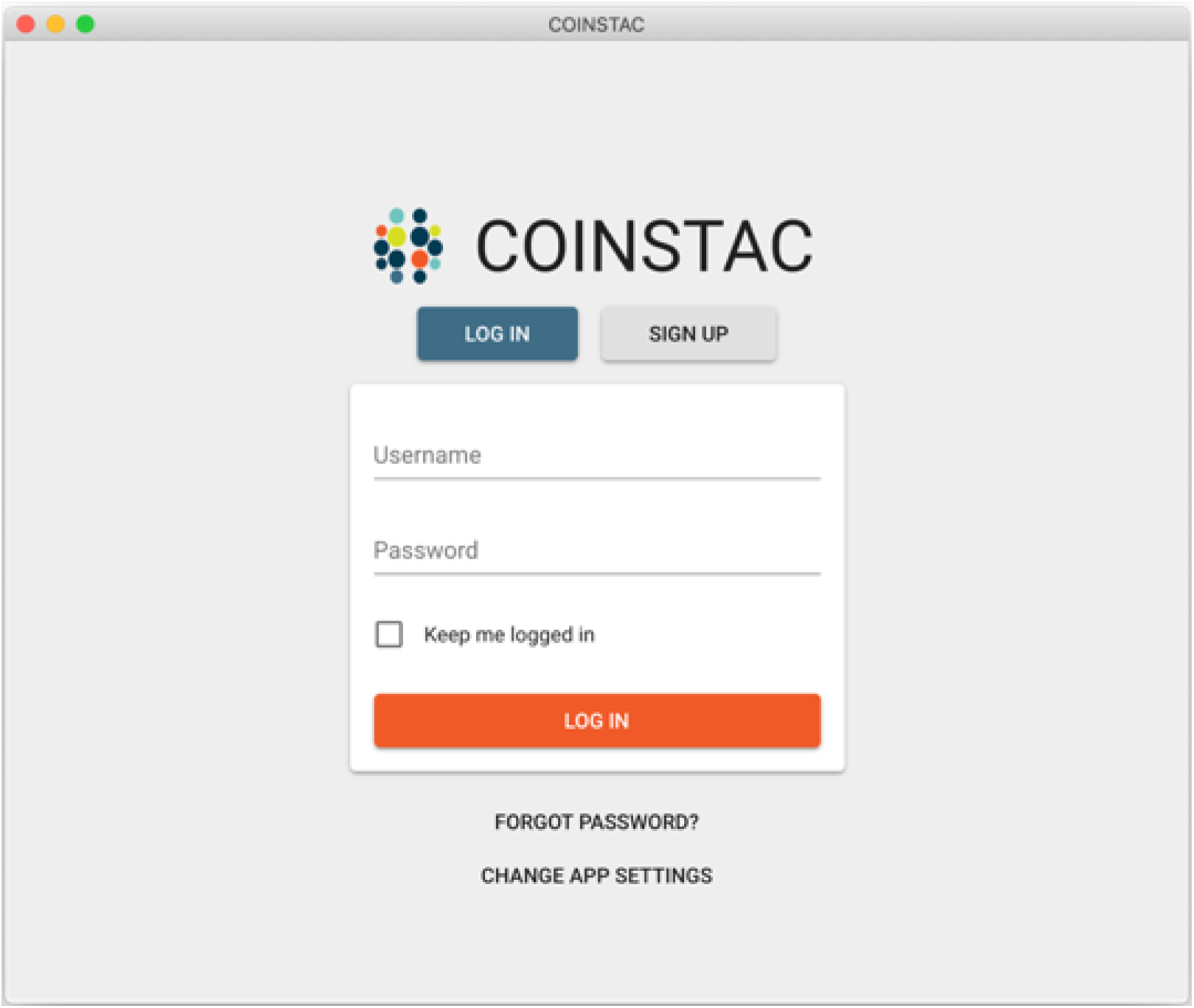
COINSTAC application login/home page.

**Figure 2:**
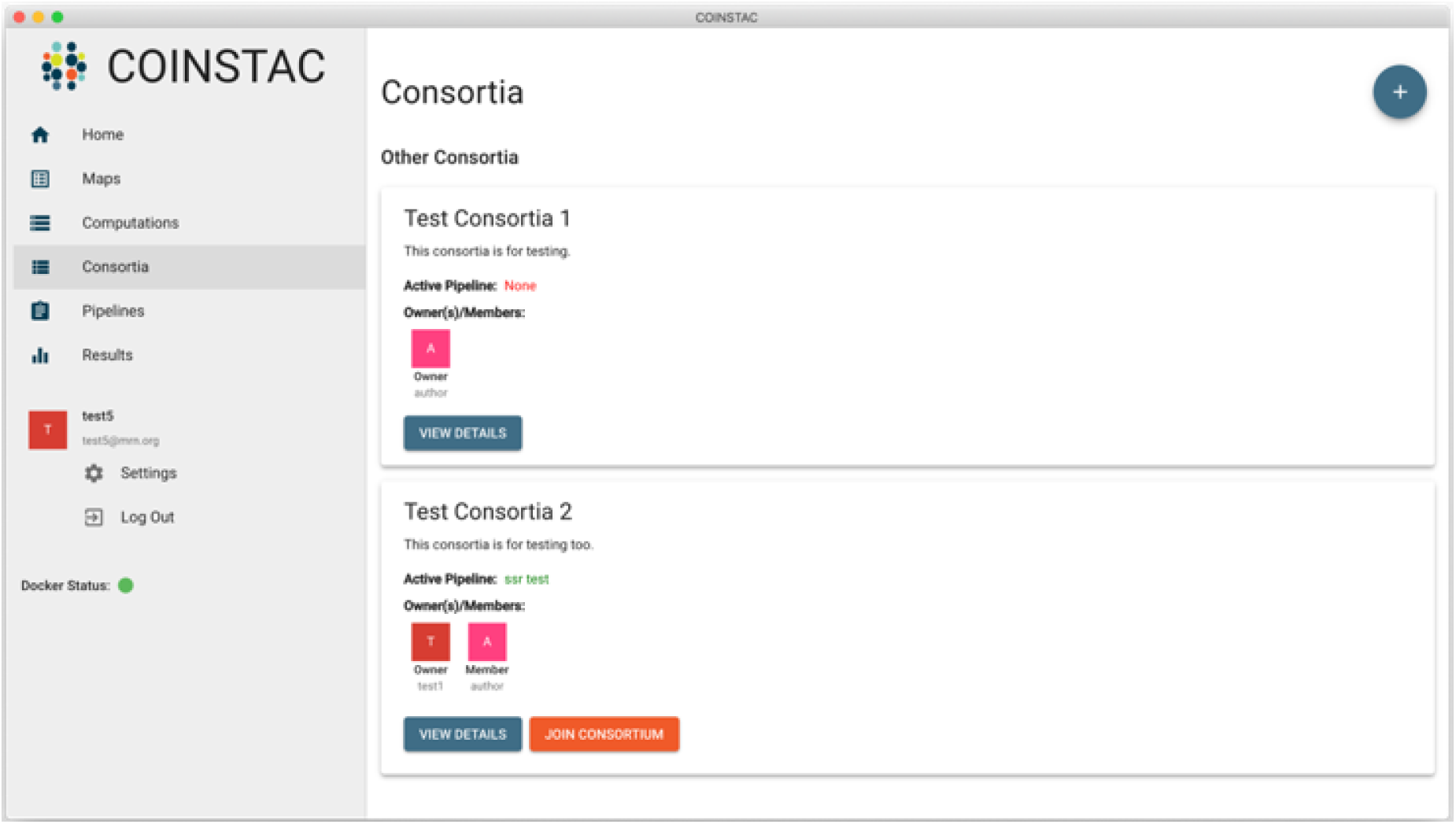
COINSTAC application Consortia page.

After a consortium has been created, the owner will be redirected to a Pipeline Creation page where he/she can define the model that will be used in the decentralized analysis. The consortium owner can choose the type of analysis (the computation) and then specify what variables are in the model (Figure 3).

**Figure 3:**
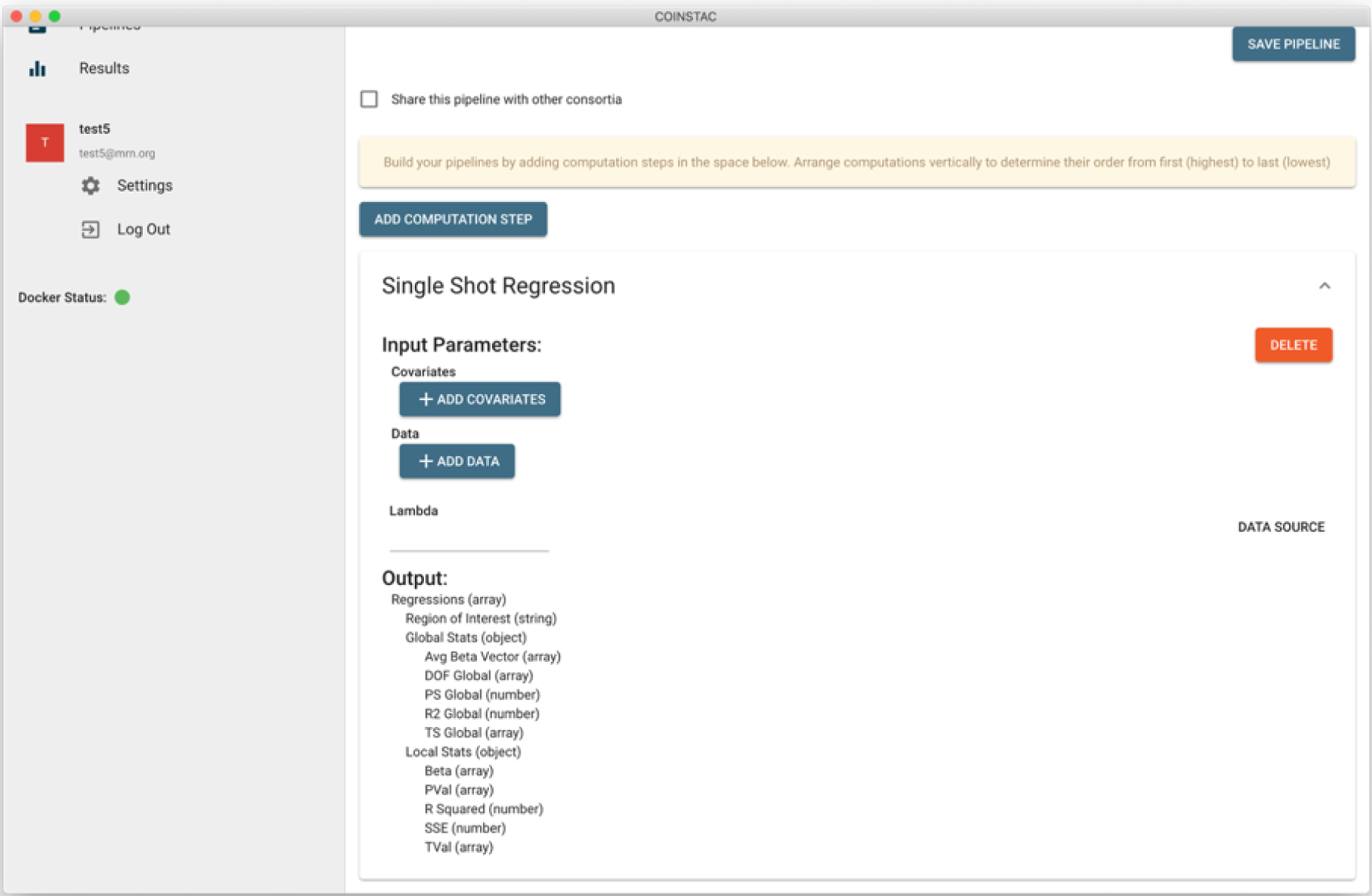
COINSTAC application Pipeline page.

Now that you the owner has created a pipeline, all members join the consortium and map their data to the variables in the model created earlier. At this point users will be taken to a Maps Page where they will be asked to point to the data to used for the analysis. When all the members are done mapping their data to the pipeline, the consortium owner is ready to start the computation.

## 3 Data

### 3.1 IMAGEN

The IMAGEN project is a longitudinal study of adolescent brain development and mental health in Europe [Schumann et al., 2010]. Participants were recruited from eight study sites in England, Ireland, France and Germany. Each study site obtained ethical approval from the local research ethics committee. Written assent and consent were acquired from all the participants and the parents prior to participation. In this research, we used structural neuroimaging, BMI and questionnaire data acquired at age 14.

#### 3.1.1 Measure of Smoking

Measures of smoking were obtained from self-reports in the Fagerstrom Test for Nicotine Dependence (FTND) questionnaire [Heatherton et al., 1991]. The answer to the question regarding lifetime smoking experience was used create a binary variable, indicating whether the participant had ever smoked (labelled as 1) or not (labelled as 0).

#### 3.1.2 Structural Magnetic Resonance Imaging

High-resolution T1-weighted structural images were acquired using 3T MRI scanners (Siemens, Philips and General Electric) across all IMAGEN study sites. An MRI sequence based on the ADNI protocols (http://adni.loni.usc.edu/methods/documents/mri-protocols/) was used. Visual inspection was performed to exclude low-quality images (movement artefacts, brace artefacts and field inhomogeneities, etc.). Grey matter volumes (GMVs) were obtained from voxel-based morphometry (VBM) analysis, by using the VBM8 toolbox (http://www.neuro.uni-jena.de/vbm/). For the VBM analysis, structural images were first segmented into grey matter, white matter and cerebrospinal fluid. The DARTEL toolbox [Ashburner, 2007] was used to convert the images to the Montreal Neurological Institute (MNI) standard space. The grey matter volumes were modulated by the Jacobian determinant obtained from the previous step, and then smoothed with an 8mm-FWHM (full width at half maximum) Gaussian kernel.

### 3.2 cVEDA

The Consortium on Vulnerability to Externalising Disorders and Addiction (cVEDA) is a multi-site, collaborative, cohort study in India with an accelerated-longitudinal design and planned missingness approach that covers an age-span of 5-24 years 1. The cohort was setup to examine neurobehavioural developmental trajectories and vulnerability to psychopathology, with a specific focus on externalising spectrum disorders. The cVEDA MRI sample comes from 6 sites with 3T MRI scanners. The T1w imaging protocol was adapted from the ADNI consortium (ADNI-2/ADNI-GO) [Beckett et al., 2015], while ensuring comparability to the European IMAGEN consortium. Participants from 3 sites (NIMHANS, SJRI, and RVRHC) scanned at the Bengaluru site at NIMHANS using two MRI scanners (Siemens Skyra and Philips Ingenia). The remaining 3 MRI sites were at Chandigarh (Siemens Verio), Mysore (Philips Ingenia) and Kolkata (Siemens Verio). In this research, we used baseline T1w structural MRI scans from 1057 participants. Refer to Table 1 for sample characteristics and Table 2 for Acquisition Parameters. Each study site obtained ethical approval from the local research ethics committee. Written assent and consent were acquired from all the participants and the parents prior to participation.

**Table 1:**
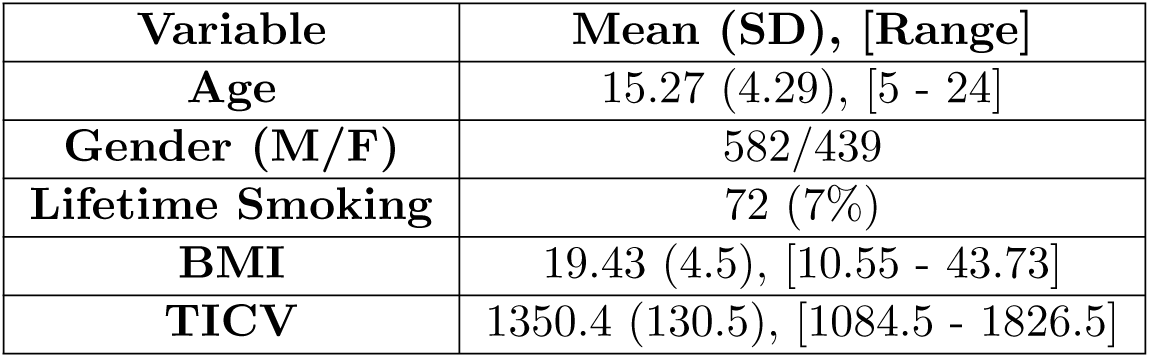
Simple Statistics from cVEDA data.

| Variable | Mean (SD), [Range] |
| --- | --- |
| Age | 15.27 (4.29), [5 - 24] |
| Gender (M/F) | 582/439 |
| Lifetime Smoking | 72 (7%) |
| BMI | 19.43 (4.5), [10.55 - 43.73] |
| TICV | 1350.4 (130.5), [1084.5 - 1826.5] |

**Table 2:** Scanner parameters from cVEDA data.

| Acq Seq | Site | Scanner Model | dx (mm) | dy (mm) | dz (mm) | TR (ms) | TE (ms) | FA* | Matrix Size (mm) | Sag Slices |
| --- | --- | --- | --- | --- | --- | --- | --- | --- | --- | --- |
| A | Mysuru | Ingena | 1.2 | 1 | 1 | 6.9 | 3.2 | 9 | 256×256 | 170 |
| B | Bengaluru | Ingenia | 1.0 | 1 | 1 | 6.5 | 2.9 | 9 | 256×256 | 211 |
| C | Bengaluru | Skyra | 1.2 | 1 | 1 | 2300 | 3 | 9 | 256×240 | 176 |
| D | Kolkata | Trio | 1.2 | 1 | 1 | 2300 | 3 | 9 | 256×240 | 176 |
| E | Chandigarh | Verio | 1.2 | 0.5 | 0.5 | 2300 | 3 | 9 | 512×240 | 176 |

#### 3.2.1 Data Preprocessing

Visual inspection was performed to exclude low-quality images (movement artefacts, brace artefacts and field inhomogeneities, etc.). AFNI’s “fat proc axialize anat” function was used for AC-PC alignment. The datasets were resampled (using a high-order sinc function to minimize smoothing) to a 1mm isotropic voxel size. Briefly, all AC-PC aligned 3D T1-weighted MRI scans were normalized using a linear affine followed by non-linear registration and corrected for bias field in homogeneities, and then segmented into GM, WM, and CSF components. We used the Diffeomorphic Anatomic Registration Through Exponentiated Lie algebra algorithm (DARTEL) to normalize the segmented scans into a standard MNI space. The segmentations were then modulated by scaling with the amount of volume changes due to spatial registration to preserve tissue volume. Finally, the modulated grey matter segmentation was smoothed at 8mm**3 FWHM Gaussian kernel 2.

Measures of smoking were obtained from WHO’s Alcohol, Smoking and Substance Involvement Screening Test (ASSIST) questionnaire that provides specific substance involvement score indicating the risk levels. A binary variable for Lifetime Smoking was created with tobacco involvement score of ≥ 3 labelled as 1 (Present) and a score *<* 3 was labelled as 0 (Absent).

## 4 Results

The use of a decentralized analytic framework like COINSTAC has many advantages. The final results are comparable to its pooled counterpart (where all the data is at one place) guaranteeing a virtual pooled analysis effect by a chain of computation and communication process. Other advantages include data privacy and support for large data

Once the two consortium members (IMAGEN and cVEDA) opened the COINSTAC application on their repective machines, joined the computation consortium and mapped the data, the consortium owner proceeds with starting the computation and it roughly took 30-45 minutes of computation time to finish the whole decentralized analysis and for the results to be displayed on the output screen as well as be downloaded for further processing. As noted earlier, a decentralized VBM was performed to understand the brain structural changes linked to BMI and smoking. Two separate models were run for the purpose of this analysis viz., age + BMI and age + smoking. Gender and site covariates were also included in both the models.

We will provide a quick summary of the results here and will discuss them further in the following section. Please note that all the maps have been thresholded with a 0.05 FDR correction [Benjamini and Hochberg, 1995]. A decentralized VBM of the structural MRI data for the effects of smoking indicates that non-smokers have higher gray matter concentration in the right anterior cingulate cortex, bilateral thalamus and brainstem, as well as the left dorsal prefrontal cortex in addition to an increase in precuneus (see Figure 4). On the other hand, for a model with BMI, Figure 5, we see increases in bilateral putamen, bilateral hypothalamus as well as lateral and medial temporal lobes on both sides. Although the age covariate was included in both the models, the effects of age for both smoking and BMI look similar and hence we report the results from only one model. From Figure 6, for age, we see widespread decreases in gray matter (weighted more towards frontal) with bilateral increases in hippocampus.

**Figure 4:**
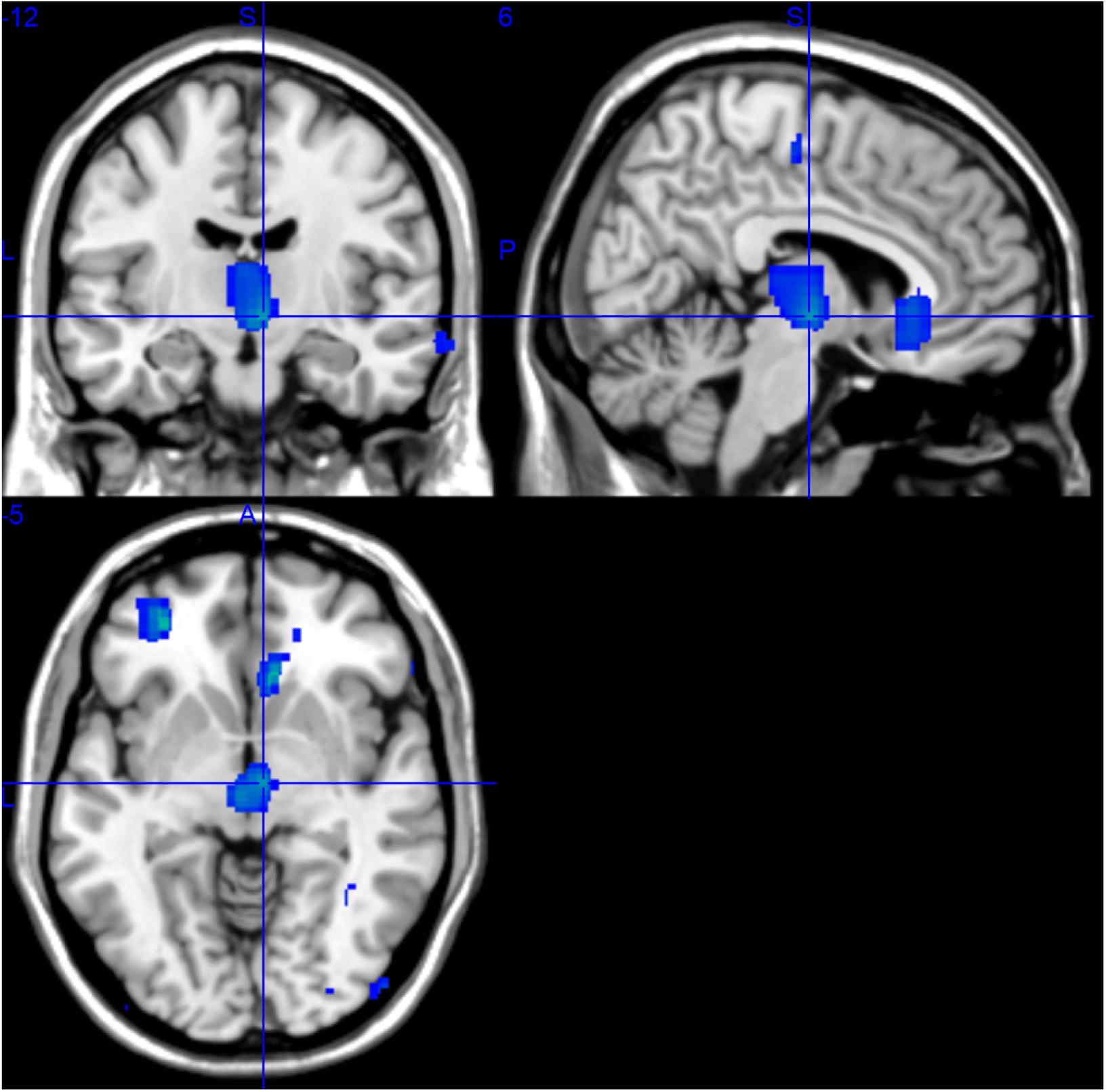
Rendered images of voxel-wise significance values (−*log*_10_*p*-value×sign(*t*)) for the covariate ‘Smoking’.

**Figure 5:**
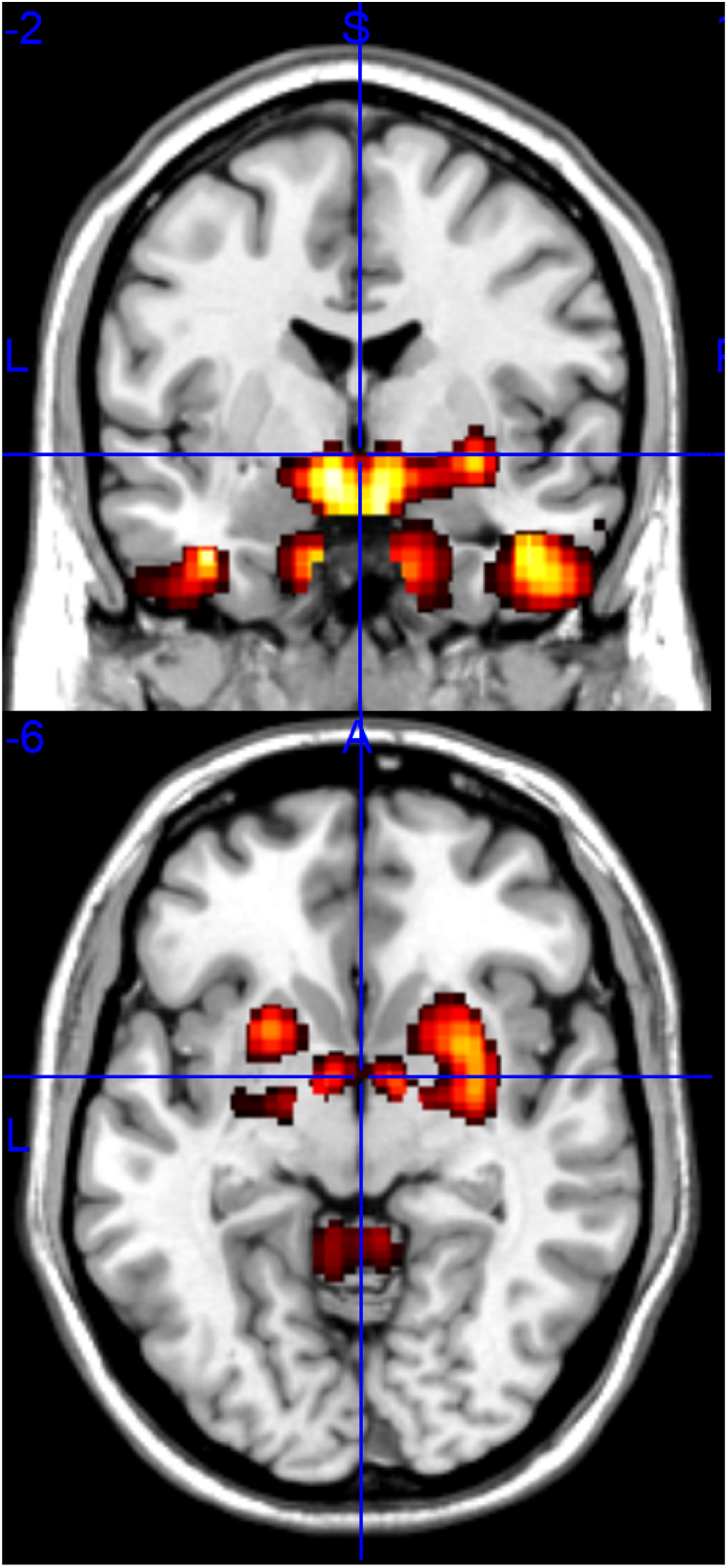
Rendered images of voxel-wise significance values (−*log*_10_*p*-value×sign(*t*)) for the covariate ‘BMI’.

**Figure 6:**
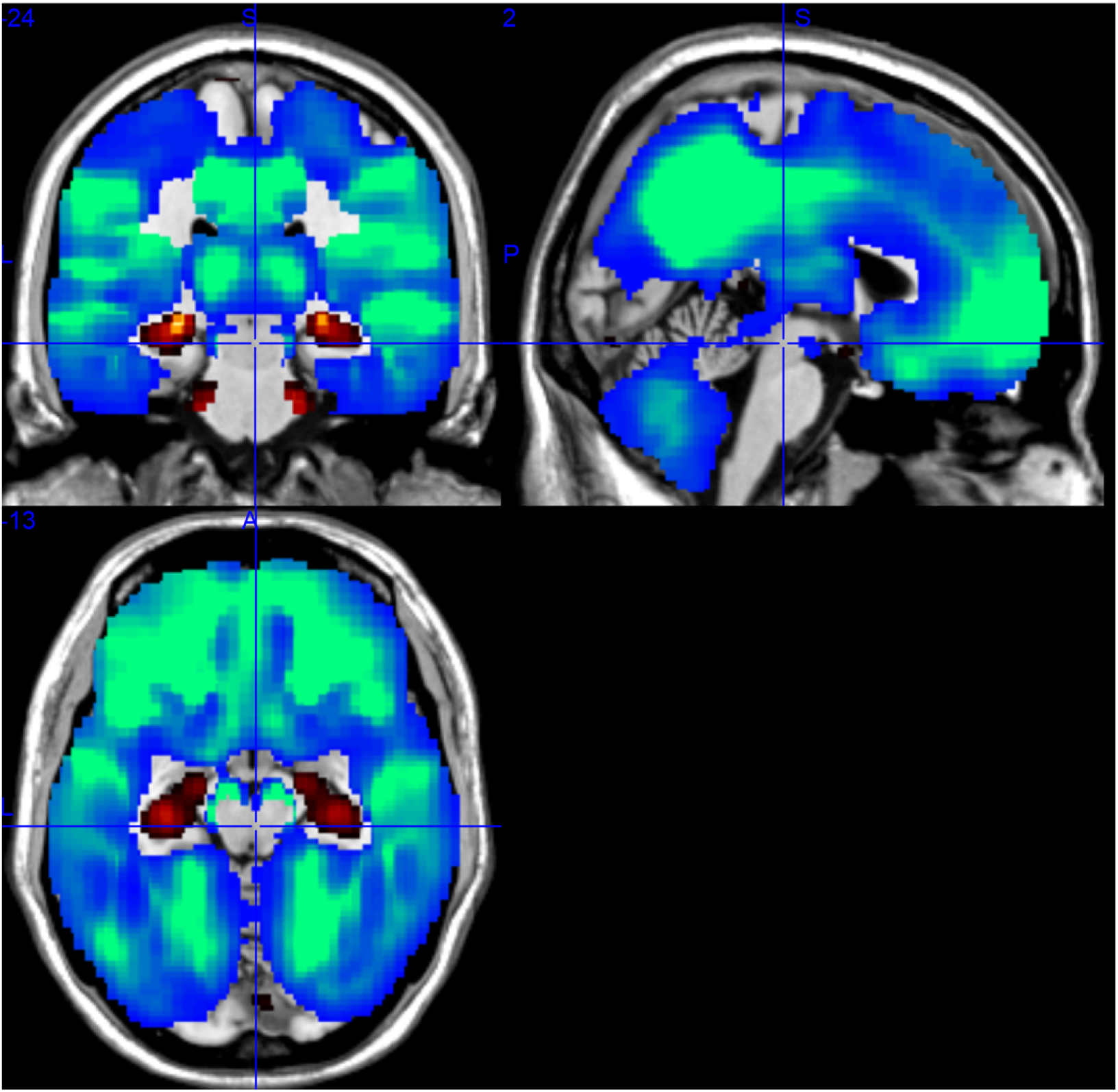
Rendered images of voxel-wise significance values (−*log*_10_*p*-value×sign(*t*)) for the covariate ‘Age’.

## 5 Discussion

The goal of this paper was two-fold: to demonstrate the feasibility of COINSTAC for performing decentralized analysis on a large data present across multiple sites and to use this experiment to understand the influence of smoking and BMI on brain structural changes. With that, we surmise this was a useful exercise working with real users of the COINSTAC application and we believe the success of this exercise culminating with this writing will spawn more fruitful collaborations going forward. Overall, in the words of the users (IMAGEN and cVEDA), the ability to examine the voxel-wise effects as against the traditional ROI-wise summary effects across multiple sites with data-privacy issues handled and with efficient decentralized computations is what they found unique and an exciting prospect for further collaborations. From a development standpoint, it was a great experience for the developers of the COINSTAC application as it gave them a unique insight into how the research conversations happen and would serve them well in easing out the user experience associated with the use of COINSTAC.

We reckon that the results in this study were consistent with existing findings from literature. In addition, they extend some findings which we will discuss here under. We noted earlier that non-smokers have higher gray matter concentration in the right anterior cingulate cortex, bilateral thalamus and brainstem, as well as the left dorsal prefrontal cortex in addition to an increase in precuneus. From literature, orbitofrontal cortex and smoking association is fairly understandable with links to impulsivity/decision making and supported by literature. However, the causal relationship is yet debatable. Olfactory cortex involvement is slightly novel given the large N in this study, but has been linked to smoking previously [Schriever et al., 2013].

While studying the influence of BMI, we reported seeing increases in bilateral putamen, bilateral hypothalamus as well as lateral and medial temporal lobes on both sides. This is an intriguing finding indicating there could be a non-linear effect. In one previous study, BMI correlated with brain activity in the left putamen, amygdala and insula in an inverted U-shaped manner [Dietrich et al., 2016]. The hypothalamic link to BMI is interesting as well, as the hypothalamic centers have a role in regulating food intake. However the reported literature points towards an atrophy rather than an increase with BMI [Kurth et al., 2013]. They report a negative correlation with BMI and waist circumference. However, the BMI range in their work was 18.18 to 42.37, thereby making it unlikely that had a problem with non-linearity of effects as they did not have low BMI group (*<* 18.5). We surmise that this effect maybe specific for adolescents, where hypothalamic volume increases with BMI and then starts reducing with chronic obesity in adulthood.

We also reported that there’s a widespread decrease in gray matter (weighted more towards frontal) with bilateral increases in hippocampus. Gray matter volume reduction in adolescence is fairly straighforward and is linked to maturation (cortical thinning/synaptic prunning/increase in underlying WM etc). However, the DMN localization of VBM-related grey matter reduction in adolescence is a fascinating insight. Maturation of the association cortices continues well into the adolescence. DMN mainly involves regions of the association cortex, and evolutionarily speaking, these regions are placed at an increased spatial distance from sensory-motor areas, the latter maturing much earlier. The “tethering hypothesis” [Buckner et al., 2013] suggests this aspect of DMN allows cognition to become more decoupled from sensory(perception)-motor(action) cycles. The hippocampal volume increase looks clear and it reaches a plateau only by mid adulthood.

## 6 Conclusion

In this paper, we showcased the power of COINSTAC in enabling neuroimaging researcher to answer important research questions by performing a multi-site study without having to pool the data. Decentralized voxel-based morphometric analysis of structural magnetic resonance imaging data from two different sites in UK and INDIA revealed some interesting insights into the gray matter concentrations in adolescent brains as a function of age, body mass index and smoking. Other advantages of such a decentralized platform include data privacy and support for large data. In conclusion, the results presented here strongly encourage the use of decentralized algorithms in large neuroimaging studies over systems that are optimized for large-scale centralized data processing.

## 7 Conflict of interest

The authors declare no conflict of interest.

## 8 Acknowledgments

This work was funded by the National Institutes of Health (grant numbers: P20GM103472/5P20RR021938, R01EB005846, 1R01DA040487) and the National Science Foundation (grant numbers: 1539067 and 1631819).

## 9 Author Contributions

HG implemented the decentralized regression algorithms on structural MRI data and the manuscript writing. BH, and ZZ contributed data to the study as well as contributed to some parts of the writing. EV contributed to performing the analysis and organizing the results. RK developed the software for the COINSTAC platform. GS is a co-investigator and has been instrumental in facilitating this multi-site study. VC led the team and formed the vision and also helped interpret the results.

